# Evidence for fungivory in *Proctotydaeus furnarius*, a brood mite of the Australian stingless bee *Tetragonula carbonaria*

**DOI:** 10.64898/2026.10.26.762071

**Authors:** Clare Price, Jasmin Li, Ana F. Pantoja, Reilly Seet, Rosalyn Gloag, Kenya E. Fernandes

**Affiliations:** School of Life and Environmental Sciences, University of Sydney, Sydney, NSW, Australia

**Keywords:** Meliponini, stingless bees, acari, mites, fungivory

## Abstract

Mites are common but poorly understood associates of stingless bee nests. The brood mite *Proctotydaeus furnarius* inhabits brood cells of *Tetragonula carbonaria* and was hypothesised to feed on fungi. We tested this hypothesis using electron microscopy, ITS amplicon sequencing, dye-based feeding trials, and brood-cell culturing. The unarmed, needle-like stylets of *P. furnarius* resembled those of fungivorous mites. In feeding trials, *P. furnarius* consumed a yeast isolate (*Zygosaccharomyces* sp.) cultured from brood cells, although also consumed crushed bee pupae. Mite presence did not detectably alter the culturable brood-cell microbial community. These findings suggest that *P. furnarius* feeds on fungi inside host nests but could shift its diet to feed on host brood according to resource availability.

## 1 Introduction

Stingless bees (Tribe Meliponini) are ecologically and economically valuable pollinators that are managed in hives for honey and crop pollination in many countries across the tropics and subtropics (Bueno et al., 2023; Grüter, 2020; Michener, 2013). Effective management and conservation of these social bees requires knowledge of both their own biology, and the organisms that share their nests. These nest associates range from parasites to beneficial symbionts, but their effects on host health are rarely fixed, since the net effect of the same association can shift between beneficial and harmful depending on density or ecological context (Rutkowski et al., 2023; Rosa-Fontana et al., 2022; Michener, 2013; Warrit & Lekprayoon, 2011; Eickwort, 1994). Mites are among the most common but least understood of these nest associates, with their role in stingless bee colonies remaining poorly characterised for all but a handful of species (Da-Costa et al., 2021; Eickwort, 1994).

Recently, the mite *Proctotydaeus (Proctotydulus) furnarius* was found living inside the brood cells of the Australian stingless bees *Tetragonula carbonaria* and *T. hockingsi* (Seeman et al., 2025). This mite enters new brood cells at the time they are being mass-provisioned (Pantoja et al. 2026). Once brood cells are sealed by workers, the mites reproduce and end up inside the pupal cocoons of developing bees, with mites being ultimately released at adult bee emergence (Pantoja et al. 2026). The genus *Proctotydaeus* includes 21 described species with diverse lifestyles, including associations with insects such as bees, beetles, grasshoppers and moths. It also includes fungal-feeding mites that maintain obligate relationships with numerous Brazilian stingless bee genera (Rosa-Fontana et al., 2022). Given this close phylogenetic relationship to known fungivorous mites, *P. furnarius* was hypothesised to feed on fungi and play a similarly beneficial role in *T. carbonaria* and *T. hockingsi* colonies (Seeman et al., 2025).

A fungivorous role would place *P. furnarius* within a broader pattern of nutritional and ecological associations between fungi and stingless bees. Fungi can provide direct nutritional benefits, including vitamins and amino acids, as well as metabolic functions such as detoxification of stored food and pollen compounds (Rutkowski et al., 2023). One Brazilian stingless bee species requires a specific fungus, identified as *Zygosaccharomyces* sp., during development to obtain sterols essential for larval pupation (Paludo et al., 2018). However, fungi can also negatively affect bee colonies, with many taxa occurring as saprotrophs or potential pathogens in stingless bee nests (Rutkowski et al., 2023; De Paula et al., 2021).

Whether a fungivorous mite is beneficial to its bee host may therefore depend on which fungi it consumes, from which part of the nest, and under what conditions.

Evidence from *P. furnarius* itself already suggests that its feeding behaviour and impact on its stingless bee hosts may be context dependent. A recent study found mixed evidence for the mite as a beneficial associate (Pantoja et al. 2026). Mites were observed to spend most of their time on brood cell walls while the developing bee was in the larval or egg stage, consistent with feeding on fungi within the cerumen. However, this behaviour shifted towards spending time on the bee itself once it reached the pupal stage. The same study found that high mite loads within pupal cocoons were associated with higher incidences of deformity (shrivelled wings, short abdomens) in emerging bees suggesting that at high abundances, or possibly under starvation conditions, *P. furnarius* can negatively impact developing bees. This may be because at high loads, mites either feed directly on the pupa itself or their feeding alters microbial conditions within the pupal cocoon in ways that impede development.

Related fungivorous mites provide further evidence that these associations can influence bee health through multiple pathways. *Neotydeolus therapeutikos,* which feeds on a fungus thought to be a bee pathogen, has been shown to reduce larval mortality rates by almost 50% (Flechtmann & de Camargo, 1971). *Proctotydaeus (Neotydeolus) alvearii* feeds on both fungi and larval food, performing cleaning tasks within brood cells that improve larval survival, and has been shown to increase larval survival specifically under insecticide stress (Rosa-Fontana et al., 2022). These examples demonstrate that fungivorous mites can benefit their bee hosts by consuming potentially harmful fungi or modifying the brood-cell environment, though they may also compete for brood resources.

In this study, we combine morphological, molecular, and experimental approaches to assess the feeding capabilities, microbial associations, and effect on the natural brood cell environment of *P. furnarius* in nests of stingless bee host *Tetragonula carbonaria*. These efforts aim to better characterize where *P. furnarius* sits on the spectrum from obligate to facultative fungivore, and how its feeding habits shift under different conditions, and therefore to understand the effect of *P. furnarius* on the health and natural microbiota of *T. carbonaria* colonies.

## 2 Materials and Methods

### 2.1 Sample collection

We sampled *Proctotydaeus furnarius* mites from managed *Tetragonula carbonaria* hives maintained on campus at The University of Sydney. Hives were opened to expose the brood comb, and small sections (approximately 5 cm^2^, comprising ∼100 brood cells) were extracted from the retreating edge and confirmed as late pupal stage by their distinctive pale semi-transparent look. Comb sections were kept in sterile petri dishes. For feeding trials, live mites were collected directly from fresh comb using a small vacuum device and transferred immediately into Eppendorf tubes without further preservation. For all other mite-based experiments, mites were collected from fresh or frozen comb sections using a fine-tip paintbrush, pins, and forceps, deposited into tubes containing 100% ethanol, and frozen at – 80 °C until processing. Comb sections not immediately processed were stored at –30 °C.

### 2.2 Scanning electron microscopy of mite mouthparts

The external mouthpart morphology of mites can provide clues as to their likely diet, with predatory mites often possessing specialised grasping or piercing structures while fungivorous mites typically lack these structures (Shatrov, 2012; Di Palma et al., 2009; Nuzzaci & Di Palma, 2002; Di Lillo et al., 2001; Hallas, 1982; Blake et al., 1978; Hislop & Jeppson, 1976). At around 250 µm body length, these features of *P. furnarius* are challenging to observe with standard light microscopy. We therefore visualized *P. furnarius* morphology using scanning electron microscopy.

We collected approximately 30 mites each into two Eppendorfs (2 mL) of 100% ethanol. We allowed mites to settle before removal of the majority of ethanol by pipette leaving a small residual volume containing the mites (∼50 µL). Approximately 2 mL of wash solution was then added and tubes inverted. This wash process was then repeated six times (total of 2 washes of 50% HMDS/50% ethanol followed by 5 washes of 100% HMDS). On the final wash, as much HMDS was removed as possible while still retaining the mites. Samples were left overnight in a fume hood, on their sides with lids open, to allow evaporation and complete drying.

Electron microscope stubs were coated with double-sided carbon tape. Tubes containing dried mites were inverted over the tape and tapped gently to release mites onto the surface. The presence of mites was confirmed by light microscopy before proceeding. Mounted mites on stubs were then coated with 15 nm gold via sputter coater and imaged using a Thermo Scientific Phenom XL Desktop Scanning Electron Microscope. A total of five mites were successfully mounted and imaged in usable orientations (two ventral, two lateral, one dorsal).

### 2.3 Detection of fungal DNA in mite bodies

If *P. furnarius* commonly consumes fungi inside the sealed brood cells of *T. carbonaria*, then we expected to detect fungal DNA from homogenized mite tissue (including mite guts). We extracted DNA from pooled sets of *P. furnarius* whole bodies collected from the brood cells of four *T. carbonaria* hives (Hives A-D; n = 3 samples; ∼150 mites each where one sample combined mites from Hives A and B, one sample comprised mites from Hive C and one sample comprised mites from Hive D).

We first processed mites through a series of washing steps to remove microbes from their exoskeletons. Mite pools in ethanol were centrifuged for 5 minutes at 14,000 rpm and the supernatant removed. Next, 100 µL of bleach solution (5% bleach in 0.1% Tween 20) was added and incubated for 30 seconds before immediate dilution with 900 µL of sterile dH_2_O. Samples were centrifuged again and the supernatant removed. To remove residual bleach, samples were washed twice with 1 mL sterile dH_2_O, vortexing briefly before each 5-minute centrifugation and removing the supernatant after each. A negative control consisting of 20 µL of the final dH_2_O wash was generated at this stage to process alongside samples. All samples were then freeze-dried at 0.3 mbar for 7 hours and stored at –80 °C until DNA extraction.

DNA was extracted using the MP Biomedicals Magbeads FastDNA Kit for Faeces, modified for mites such that the initial kit wash steps were removed and freeze-dried samples were instead homogenised directly at 6,000 rpm for 40 seconds twice, with a 2-minute break in between. Extracted DNA was tested via PCR and gel electrophoresis to confirm the presence of fungal DNA. Samples were submitted to the Ramaciotti Centre for Genomics (University of New South Wales, Sydney) for paired-end amplicon sequencing on the Illumina MiSeq v3 platform (2 x 300 bp) using the ITS1F-ITS2 primer pair (Smith & Peay, 2014) for the ITS1 region.

We analyzed ITS1 sequence data using the DADA2 pipeline in RStudio (version 4.4.0). Reads were filtered with a minimum length of 50 bp and maximum of two expected errors, followed by standard DADA2 denoising, paired-end merging, and chimera removal. Amplicon sequence variants (ASVs) were taxonomically assigned using the UNITE general FASTA v10.0 for ITS (Abarenkov et al., 2024). Unclassified ASVs were additionally queried against NCBI’s nucleotide database using BLASTn; ASVs with no matches, or that matched bacterial or bacteriophage sequences, were considered non-fungal artifacts and removed from the dataset. After processing, there was an average of 18,745 ± 1,214 reads (mean ± SEM) per pooled mite sample and 5,014 reads in the negative H_2_O control. ASVs present in the negative control were subtracted from all samples (3/10 ASVs; 1.1% of total test sample reads).

### 2.4 Feeding trials

To assess the capacity of *P. furnarius* to feed on different resources available inside the brood cells of their host, we incubated live mites together with either microbial cultures or brood comb components that had been dyed blue. As the mite bodies are translucent, any blue dye visible in the digestive tracts of mites would indicate consumption.

To ensure that we were offering mites potential food sources that were naturally present in the brood cell environments they inhabit, we first cultured and isolated bacterial and fungal colonies from *T. carbonaria* brood comb cells. Individual brood cells containing mites (n = 3) were extracted from frozen brood comb (–30 °C overnight) using sterile scalpels and forceps. Cells were opened and pupal bees removed before depositing cells in 2 mL of one of three buffers (0.1% peptone, PBS or 0.01% Tween 20), vortexing, and incubating for at least 5 minutes at room temperature. From each sample, 25 µL was plated onto 4 agar types in duplicate: PDA, NA, 15GF and MYAG50, and incubated at either 25 °C or 30 °C for 4 weeks (n = 24 plates total). Morphologically distinct colonies were isolated and purified by re-streaking on the same media type. Purified colonies were identified by Sanger sequencing of the ITS1 or 16S (V3-V4) regions using the PCR methods described above, with sequencing performed by the Australian Genome Research Facility. Resulting sequences were identified to their closest sequenced match using NCBI BLASTn (Supplementary Table S1). A total of 62 isolates were identified, yielding a total of 19 unique genera.

We then selected nine microbial isolates for use in feeding trials: four fungal (*Zygosaccharomyces* sp., *Cystobasidium* sp., *Monascus* sp., *Cladosporium* sp.) and five bacterial (*Stenotrophomonas* sp., *Roseomonas* sp., *Novosphingobium* sp., *Pseudomonas* sp., *Rhizobium* sp.) isolates. Selected identified microbial isolates were cultured on their required media, then scraped into 500 µL sterile dH_2_O. We also selected four bee brood components for feeding trials: brood cocoon (with bee removed), late-stage pupae, larvae, and larval food. All brood components (excluding larval food) were obtained from frozen comb sections and washed with 0.4% bleach in 0.04% Tween to remove excess material and microbes, then washed twice with sterile dH^2^O and crushed in 500 µL sterile dH_2_O using micro pestles. Finally, we included two controls in feeding trials: PDA media scraped into sterile dH_2_O (to account for the possibility that mites were feeding on residual PDA used to culture the microbe, rather than the microbe itself) and sterile dH_2_O only (to account for some microbes and brood components being suspended in water).

We dyed solid foods by adding a 500 µL aliquot of food-grade blue dye (final concentration 50%), incubating for 2 minutes, then centrifuging for 1 minute at 14,000 rpm. The dyed supernatant was removed and the pellet washed twice with 1 mL sterile dH_2_O. Solid foods included *Zygosaccharomyces* sp., *Cystobasidium* sp., *Monascus* sp., *Cladosporium* sp., *Roseomonas* sp., *Pseudomonas* sp., brood cocoon (with bee removed), late-stage pupae, larvae, and the PDA control. We dyed liquid foods directly, with no wash performed. Liquid foods included the *Novosphingobium, Stenotrophomonas,* and *Rhizobium* cultures, which formed suspensions, larval food, and the water control.

We performed three trials per treatment, with ∼ 300 mites per treatment (an estimate, given the small size and delicate bodies of mites). A total of 4,731 mites were used across the 45 trials (4,480 following exclusion of mites found on or within food), with an average of 99.6 mites per trial (range 31-213). Mites were starved in the dark for 24 hours at room temperature prior to trials. For each trial, a ∼5 mm^2^ drop or dollop of dyed food was placed at the edge of a 35 mm petri dish, starved mites tipped onto the opposite side, and the dish sealed with parafilm. Mites were incubated with food in the dark for >7 days at room temperature, by which point all individuals were deceased. Plates were then examined under a dissection microscope and mites classified as dyed (visible blue colouration) or undyed (no blue visible). Mites found on or in the food were counted but not included in analyses as their dyed status could not be reliably determined. Each trial was conducted with three independent replicates, and the percentage of dyed mites for each food was compared against the water control using Fisher’s exact test, with p-values adjusted using the Benjamini-Hochberg method.

To confirm consumption during feeding trials at the molecular level, we performed an additional three food trials for one fungal treatment (*Zygosaccharomyces sp.*), a whole crushed brood cell (including bee pupae), and no food (starvation). For these additional trials, we then performed whole-body DNA extraction and ITS1 amplicon sequencing on deceased mites (n > 50 per trial) which were washed out from food using 100% ethanol. We then performed DNA extraction, sequencing and bioinformatic analysis as described above, except that a *Zygosaccharomyces sp.* isolate cultured from *T. carbonaria* brood cells and identified via Sanger sequencing was additionally used as a query against all ITS ASVs (BLASTn package in R, at a 95% identity cutoff). This allowed us to confirm that the specific *Zygosaccharomyces* ASV fed to the mites was present in their bodies.

To test directly whether *P. furnarius* could pierce the intact cuticle of bee pupae, we additionally attempted trypan blue vital-staining assays and whole-pupa dye immersion, following approaches developed for *Varroa destructor* (Kanbar & Engels, 2004). These attempts were unsuccessful due to the comparatively small size of *P. furnarius* relative to *V. destructor*.

### 2.5 Brood cell microbial community profiling

To test whether *P. furnarius* affects the culturable microbial community of the brood cells it inhabits, we compared microbial communities cultured from brood cells with and without visible mites, across four media types and hives.

Individual brood cells from three hives were classified using light microscopy as containing mites (high) or no visible mites (low) then immediately submerged in 2 mL of 0.1% peptone buffer and incubated for at least 10 minutes at room temperature. Each hive contributed 3 high- and 3 low-mite samples (n=18). From each sample, 50 µL was plated onto 4 agar types: Potato Dextrose Agar (PDA; Oxoid), Nutrient Agar (NA; Oxoid), 15% Gluctose Fructose Agar (15GF; Paludo et al. 2018), and 50% Glucose Malt-Yeast Agar (MYAG50; Detry et al. 2020). Plates were incubated at 30 °C for 4 weeks, and colonies quantified by count and number of distinct morphological types per plate.

Community composition was analysed in R using PERMANOVA, PERMDISP, and MVABUND. A single outlier plate (colony count >300, versus ≤50 for all others) was excluded from reported statistics; this exclusion was checked against the full dataset to confirm it did not substantially alter results. PERMANOVA and PERMDISP required additional exclusion of plates with zero colony growth (11 of the remaining 71).

## 3 Results

### 3.1 Scanning electron microscopy of mite mouthparts

The external mouthpart morphology of *P. furnarius* is consistent with a fungivorous diet. Predatory and parasitic mites typically bear specialised grasping or piercing structures on their feeding appendages, such as claws, hooks, or teeth. *P. furnarius* lacked these features entirely. Its paired sensory appendages (pedipalps), positioned either side of the other mouthpart region (gnathosoma), lacked claws or coarse bristles at their tips (Fig. 1A-B). Its core feeding structures (chelicerae) terminate in a pair of needle-like stylets, enclosed by lateral folds of cuticle (lateral lips), approximately 3-4 µm long, straight, and only slightly curved at their tips (Fig. 1B-C). This smooth, unarmed stylet morphology resembles that of known fungal-feeding (fungivorous) and plant-feeding (phytophagous) mites, and is distinct from the hooked, toothed structures typical of predatory or parasitic mites. SEM images also showed the mite’s abdomen (idiosoma) with pronounced folding, likely a result of sample drying and indicative of a soft exoskeleton (Fig. 1A). The cuticle surface showed fine fingerprint-like ridges (striae) and numerous small bristles (setae), in addition to larger bristles visible at whole-body scale (Fig. 1B).

**Figure 1.**
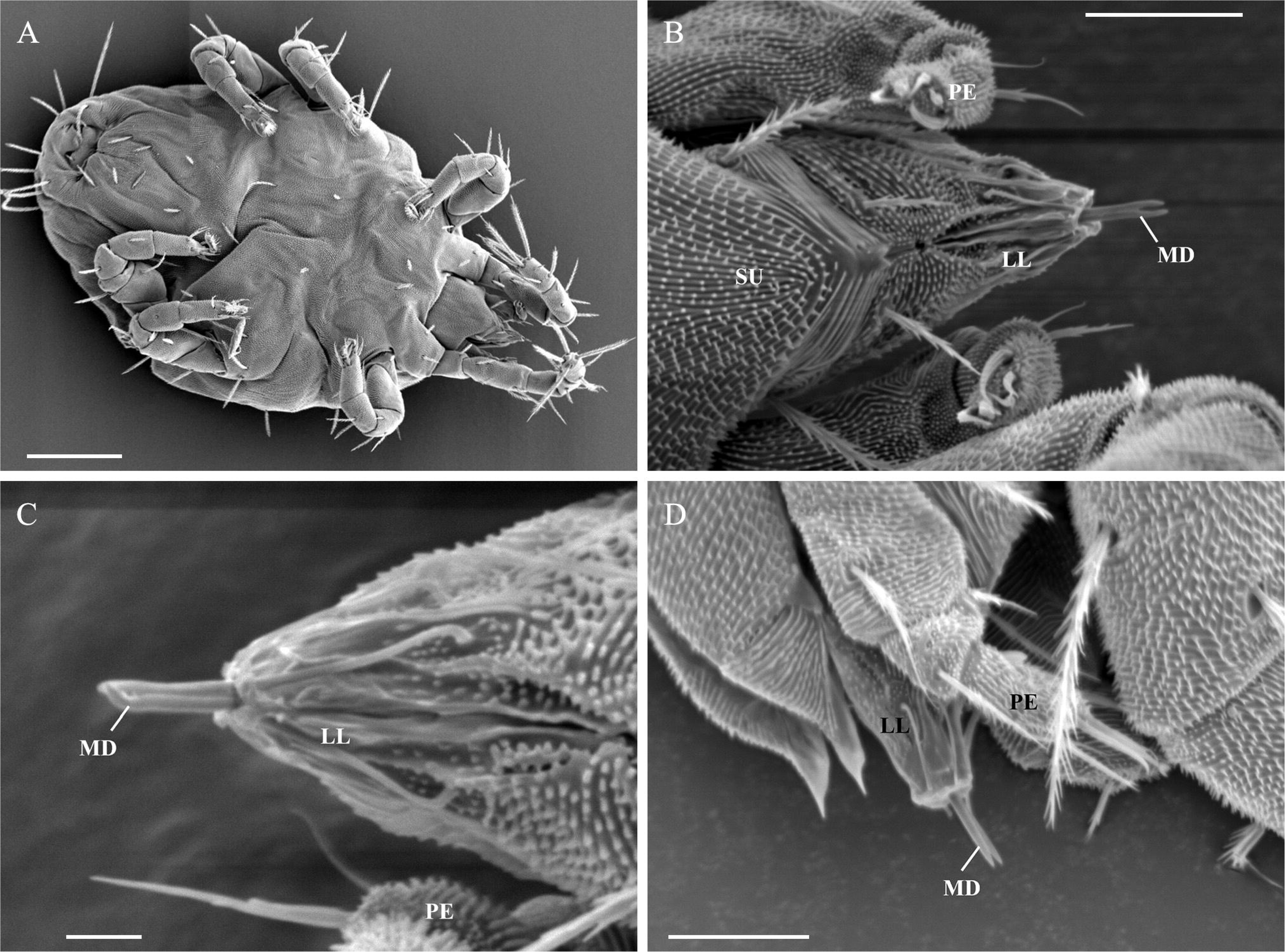
Mouthpart morphology of *Proctotydaeus furnarius*. Scanning electron micrographs showing **(A)** whole body, ventral view (scale bar = 50 µm), **(B-C)** gnathosoma, ventral view (scale bars = 10 µm (B) and 2 µm (C)), and **(D)** gnathosoma, lateral view (scale bar = 10 µm). Abbreviations: LL, lateral lips; MD, movable digits; PE, pedipalps; SU, subcaptiulum.

### 3.2 Detection of fungal DNA in mite bodies

The fungal communities within *P. furnarius* bodies were dominated by a single taxonomically unresolved amplicon sequence variant (ASV). This single ASV could not be classified beyond the kingdom level and comprised 91-99% of relative abundance across all samples (Fig. 2A). The remaining classified genera were detected at low relative absolute read counts (17-1,414 reads per sample; 150 mites per sample) and varied inconsistently between hives (Fig. 2B). *Aspergillus* and two unclassified Ascomycota ASVs were present in the sample from Hives A/B, *Malassezia* in the Hive C sample, and Cordycipitaceae and *Penicillium* in the Hive D sample.

**Figure 2.**
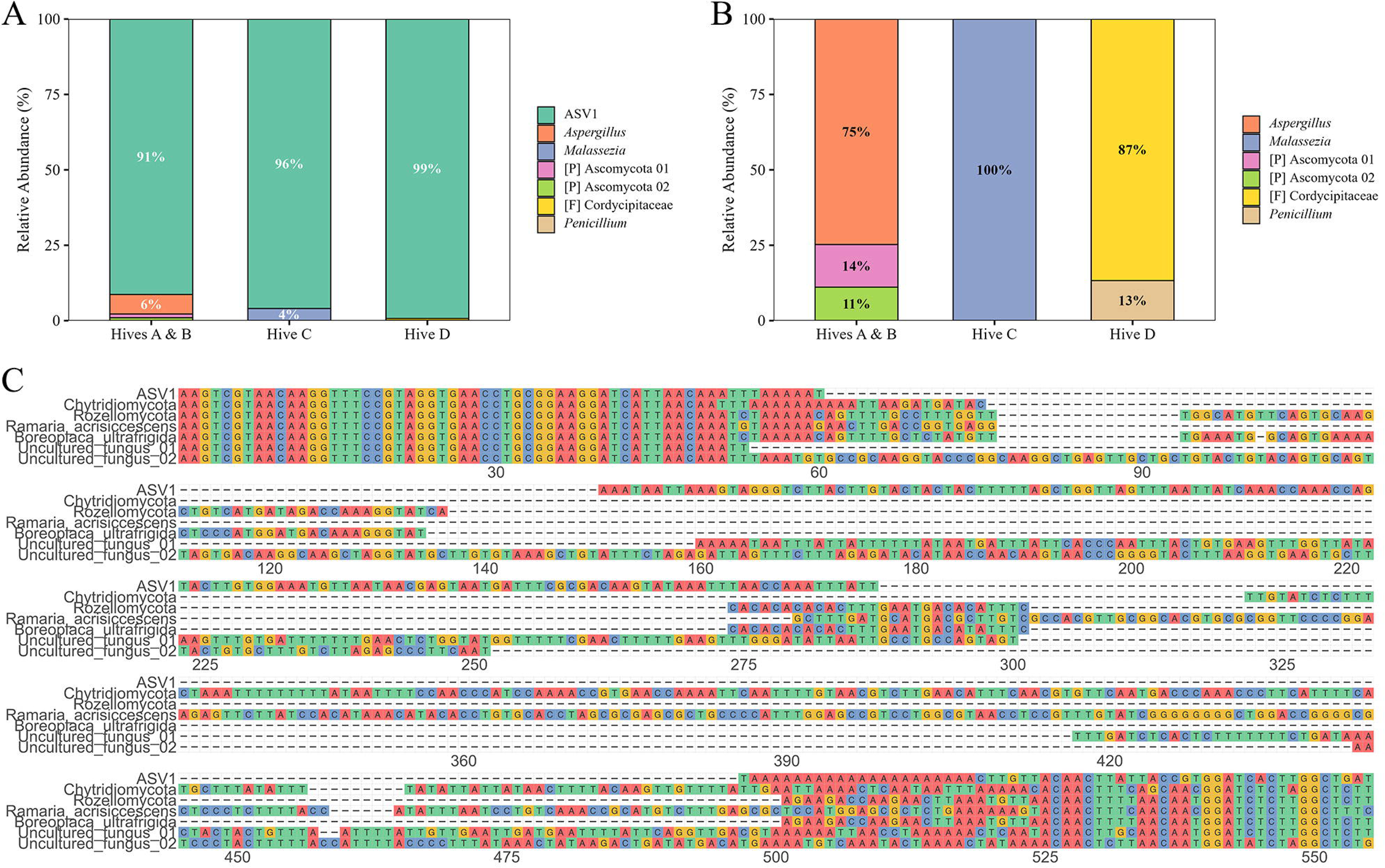
Fungal communities associated with *Proctotydaeus furnarius*. Relative abundance of fungal taxa from whole-body ITS1 amplicon sequencing of three samples (∼150 mites each) from four *T. carbonaria* hives (Hives A-D). **(A)** Full community composition, including the dominant unresolved sequence (ASV1). [P] and [F] denote taxa only resolved to phylum or family level, respectively. **(B)** Composition of the non-dominant fraction (all taxa excluding ASV1), renormalised to 100%. **(C)** Alignment of ASV1 against selected top BLAST hits, showing conserved flanking regions bordering a divergent core. Accession numbers and full hit statistics are reported in Supplementary Table S2.

The dominant unclassified ASV (“ASV1”) was queried using NCBI nucleotide BLAST, yielding a consistently low query coverage (∼25-35%) across all top hits (Supplementary Table S2). The closest matches spanned several taxonomically disparate fungal lineages, including early-diverging Chytridiomycota and Rozellomycota, the basidiomycete *Ramaria acrisiccescens,* the lichen-forming fungus *Boreoplaca ultrafrigida,* and multiple uncultured fungal clones. Alignment of ASV1 against these top hits showed conserved sequence restricted to both flanks, consistent with broadly conserved fungal ribosomal architecture rather than taxon-specific similarity, while the central region remained divergent from all hits (Fig. 2C). This pattern indicates that ASV1 likely represents a divergent, taxonomically unresolved fungal lineage.

### 3.3 Feeding trials

Dye colouration provided a reliable proxy for food consumption. Fig 3A shows the appearance of undyed mites under the microscope, and Fig 3B-C shows the appearance of dyed mites with diffuse colouring throughout their bodies. The dyed water control produced no coloured mites across three replicate trials, confirming that colouration in other trials was due to consumption of the provided food rather than of the water or dye itself.

**Figure 3.**
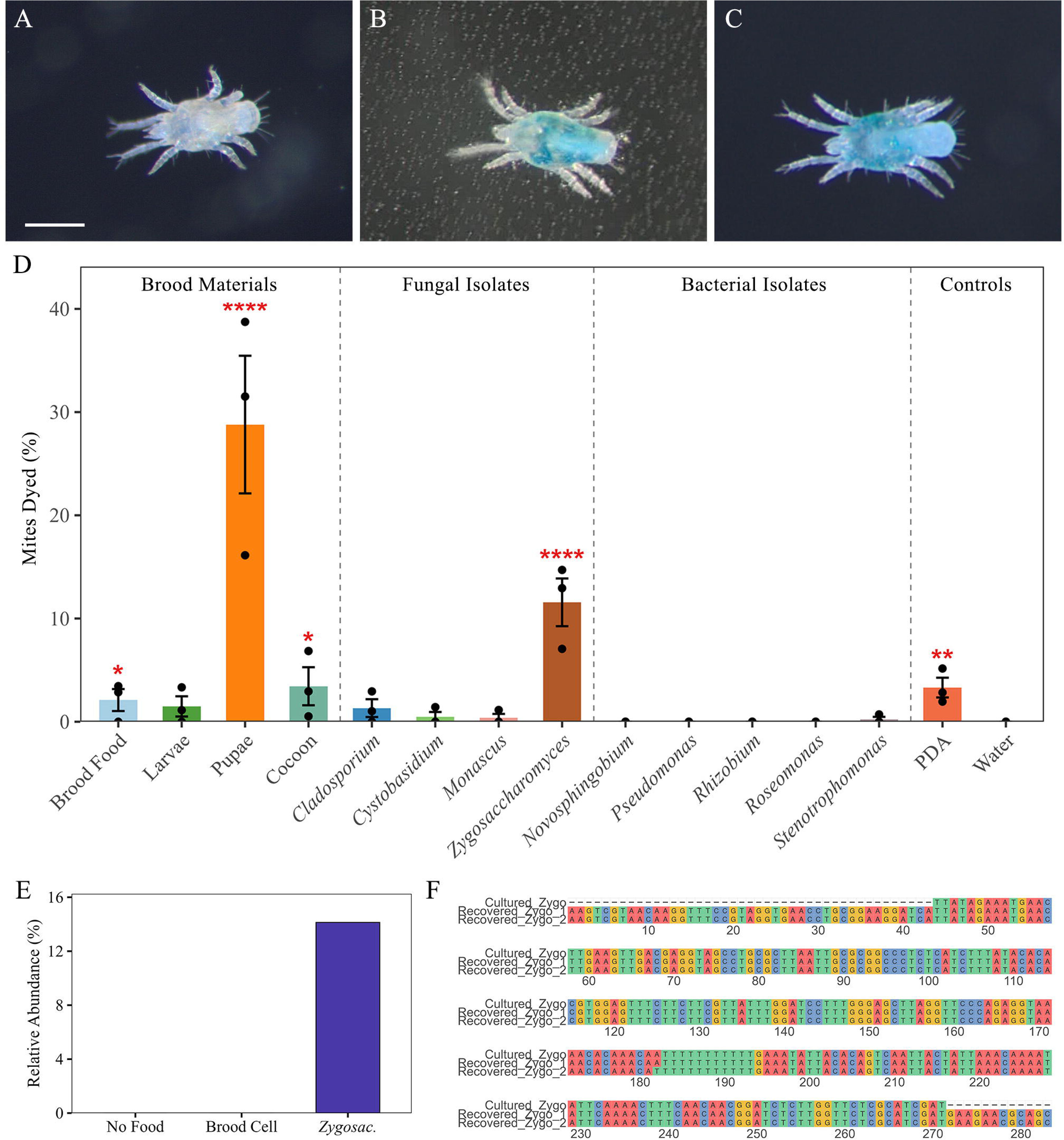
Feeding responses of *Proctotydaeus furnarius* to brood-associated resources. Representative photographs of mites from the feeding trials: **(A)** undyed mite with no visible colouration; dyed mites in **(B)** dorsal and **(C)** ventral view, showing diffuse dye colouration throughout the body (scale bar = 150 µm). **(D)** Mean percentage of dyed mites per food, grouped by brood materials, fungal isolates, bacterial isolates, and controls (n = 31-213 mites per trial; 3 trials per treatment). Points show individual replicate trials (n = 3 per food) and error bars show SEM. Red asterisks indicate statistically significant differences from the water control (p<0.05 = *, p<0.01 = **, p<0.001 = ***, and p<0.0001 = ****). Statistical results are reported in full in Table 1. **(E)** Relative abundance of the *Zygosaccharomyces*-matching ASV in mites after feeding trials with no food (starvation), a whole crushed brood cell, or *Zygosaccharomyces* sp. **(F)** Alignment of the *Zygosaccharomyces*-matching sequences from whole-body DNA extractions (Recovered Zygo 1 and 2) with cultured *Zygosaccharomyces* isolate sequence obtained via Sanger sequencing of the ITS1 region.

**Table 1.** Summary of statistical significance for each food trial relative to the water control, and sample size of each treatment.

| Food | Fisher's exact<br>test p-value<sup>1</sup> | Number of mites |  |  |
| --- | --- | --- | --- | --- |
|  |  | Undyed | Dyed | Total |
| Larvae | 0.1344 | 203 | 3 | 206 |
| Pupae | **<0.0001** | 194 | 94 | 288 |
| Brood Food | **0.0210** | 252 | 6 | 258 |
| Cocoon | **0.0192** | 290 | 7 | 297 |
| *Cladosporium* | 0.1376 | 222 | 3 | 225 |
| *Cystobasidium* | 0.8327 | 327 | 2 | 329 |
| *Monascus* | 1.0000 | 325 | 1 | 326 |
| *Zygosaccharomyces* | **<0.0001** | 211 | 27 | 238 |
| *Novosphingobium* | 1.0000 | 409 | 0 | 409 |
| *Pseudomonas* | 1.0000 | 240 | 0 | 240 |
| *Rhizobium* | 1.0000 | 268 | 0 | 268 |
| *Roseomonas* | 1.0000 | 251 | 0 | 251 |
| *Stenotrophomonas* | 1.0000 | 405 | 1 | 406 |
| PDA | **0.0037** | 415 | 13 | 428 |
| Water |  | 311 | 0 | 311 |

<sup>1</sup> P-values in bold are statistically significant.

Bee pupae produced the highest percentage of dyed mites across all three replicates (16-39%), followed by the fungus *Zygosaccharomyces* sp. (7-15%), both significantly greater than the water control (p<0.0001; Fig. 3D; all sample sizes and statistics in Table 1). The remaining brood food components all produced dyed mites (brood food 0-3%, larvae 0-3%, cocoon 1-7%), though only brood food (p=0.0210) and cocoon (p=0.0192) reached statistical significance compared to the water control. The remaining fungal isolates also all produced dyed mites (*Cladosporium* 0-3%, *Cystobasidium* 0-1%, *Monascus* 0-1%) but none reached statistical significance. As *Cladosporium* and *Monascus* could not be effectively separated from their PDA medium, a PDA control was included which was also found to produce dyed mites (2-5%) at higher proportions than the water control (p=0.0037). Of the five bacterial isolates tested, *Novosphingobium, Pseudomonas, Rhizobium, and Roseomonas* produced no visible dyed mites, and *Stenotrophomonas* produced a very small amount (0-1%; Table 1).

To confirm consumption of *Zygosaccharomyces* at the molecular level, whole-body DNA extraction and amplicon sequencing were performed on mites following feeding trials with no food, a whole crushed brood cell, and *Zygosaccharomyces*. A *Zygosaccharomyces*-matching ASV was detected at negligible relative abundance in the mites from the no food and brood cell trials but comprised 14% relative abundance in mites from the *Zygosaccharomyces* trial (Fig. 3E). This ASV was confirmed by direct comparison against the Sanger-sequenced cultured isolate used in feeding trials to match at 100% identity, providing molecular support for fungal consumption (Fig. 3F).

### 3.4 Effect of mites on the microbial community of brood cells

Mite presence had no detectable effect on the culturable microbial community of brood cells. Total colony counts varied by hive and media type, with Hive D showing consistently higher colony numbers than Hives A or B across all media, and particularly on 15GF and MYAG50 (Fig. 4A). This did not correspond to greater morphological diversity, however, as the number of colony types per plate was relatively consistent overall with almost all plates showing only one or two (Fig. 4B). Colony morphotype presence showed no consistent association with mite abundance across plates (Supplementary Table S3). Mite abundance (high or low) had no statistically significant effect on microbial community composition, with similar dispersion between mite groups (PERMANOVA, PERMDISP, MVABUND; all p≥0.46; Table 2), and this held even after accounting for hive and media type. Hive identity was the only significant factor (PERMANOVA p=0.001), driven primarily by differences in colony counts rather than colony type diversity, with elevated and more variable colony numbers in Hive D (PERMDISP p=0.005).

**Figure 4.**
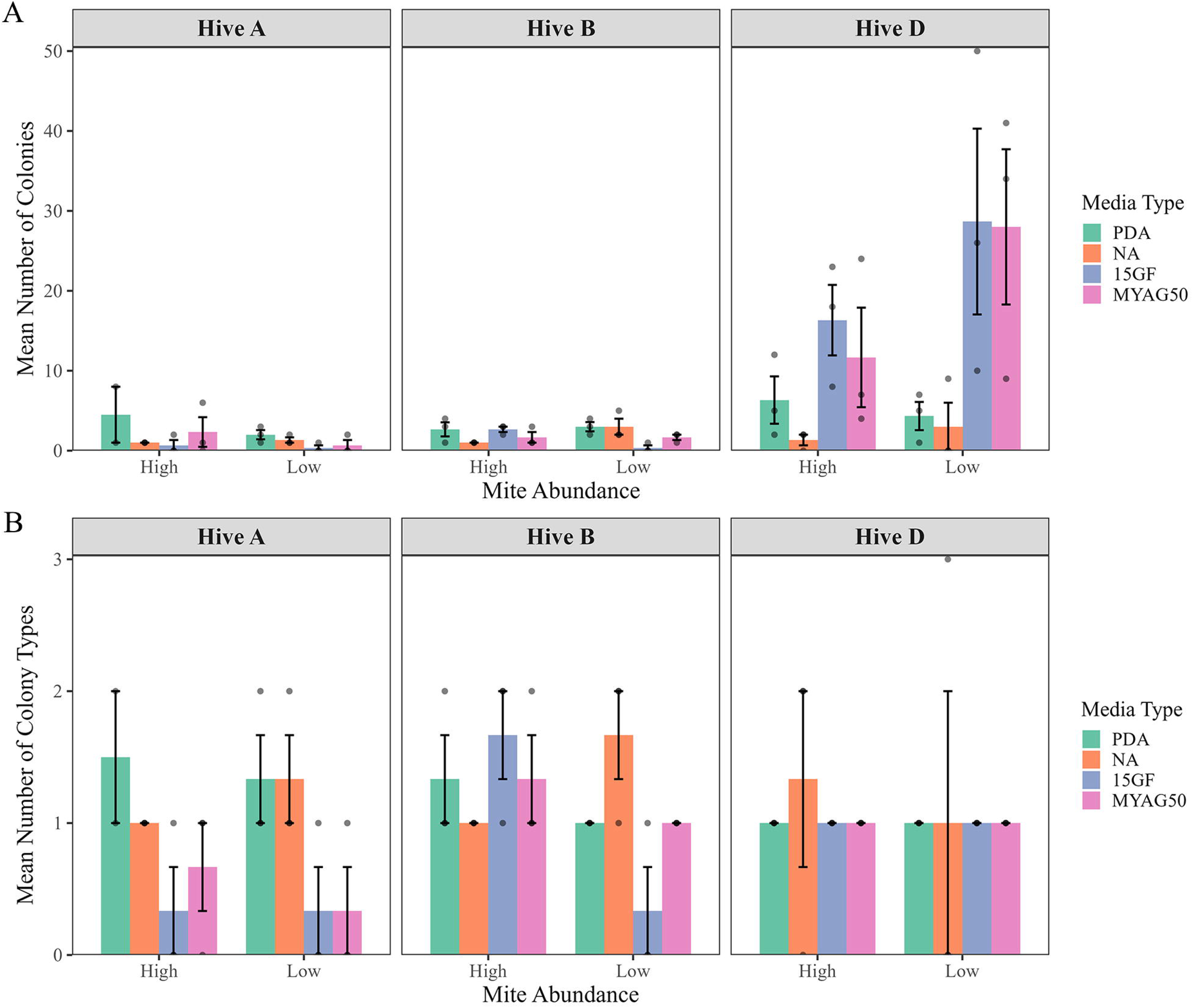
Culturable microbial communities in brood cells with high and low mite abundance. **(A)** Mean number of colonies per plate and **(B)** mean number of distinct colony morphological types per plate. Both panels are grouped by hive (A, B, D), stratified by mite abundance (High, Low) and coloured by media type (PDA, NA, 15GF, MYAG50). Points show individual brood cells (n =3 per group), error bars show SEM, and darker points indicate overlapping values. Statistical results are reported in Table 2.

**Table 2.** Summary of statistical significance for the effect of hive, mite abundance, and media type on cultural microbial community variation. ^1^.

| **Factor** | **PERMANOVA** | **PERMDISP** | **MVABUND<sup>2</sup>** |
| --- | --- | --- | --- |
| Hive | $R^2 = 0.301$<br>**P = 0.001** | **P = 0.005** | Dev = 46.85<br>**P = 0.001** |
| Mite Abundance | $R^2 = 0.008$<br>P = 0.460 | P = 0.856 | Dev = 0.52<br>P = 0.682 |
| Media Type | $R^2 = 0.071$<br>P = 0.067 | **P = 0.028** | Dev = 8.57<br>P = 0.118 |
| Hive & Mite Abundance | $R^2 = 0.025$<br>P = 0.264 | | Dev = 3.60<br>P = 0.352 |
| Mite Abundance & Media Type | $R^2 = 0.056$<br>P = 0.121 | | Dev = 4.11<br>P = 0.502 |

<sup>1</sup> PERMANOVA and PERMDISP tests required exclusion of plates with zero colony growth (11/71).
<sup>2</sup> Dev = Deviance.

## 4 Discussion

This study set out to characterise the feeding behaviour of *Proctotydaeus furnarius* and its relationship with its host, the Australian native stingless bee *Tetragonula carbonaria.* Morphological, molecular, and experimental evidence support a capacity for fungivory, consistent with the mite’s phylogenetic placement near other fungal-feeding lineages (Seeman et al., 2025; Rosa-Fontana, et al., 2022). However, feeding trials also demonstrated *P. furnarius* is capable of consuming developing bee tissue, indicating its relationship with *T. carbonaria* is unlikely to sit on a fixed point on the mutualist-to-parasite spectrum, but rather shifts with the resources available.

Feeding trials confirmed that *P. furnarius* can consume a range of dyed foods under experimental conditions. These results are insufficient on their own to characterise the complete natural diet of *P. furnarius* in brood cells, but are consistent with a broad, opportunistic diet, mirroring many other social insect-associated mites that act as fungivores, detritivores, or scavengers within host nests, often functioning as a nest “cleaning crew” that benefits the host (Okabe, 2013; Biani et al., 2009). Most notably, *P. furnarius* consumed the cultured fungus *Zygosaccharomyces*, a fungus previously identified as an important symbiont of some stingless bees (De Paula et al., 2021; Paludo et al., 2018), supported by molecular detection of a matching ASV in fed mites. This points to a plausible role for *P. furnarius* in regulating fungal symbiont populations within the nest and guarding against dysbiosis, a role that may be particularly important when hives are under stress, such as from pesticide exposure (Rosa-Fontana et al., 2022).

The reverse, however, is also plausible: overconsumption of symbiotic fungi that bees themselves depend on, particularly at high mite loads, could carry costs to the developing bee brood. This possibility would be consistent with prior observations linking elevated *P. furnarius* loads per brood cell to deformity in emerging *T. carbonaria* callows (Pantoja et al. 2026). In the present study, our parallel experiment testing whether mites measurably affected the brood cell’s culturable microbial community found no detectable effect of high mite loads, with hive identity the only significant driver of variation. Yet it remains unclear whether this null result reflects a genuine absence of effect, or simply the limited sensitivity of culture-based methods to detect modest or highly targeted symbiont interactions that are nevertheless critical to bee development.

Feeding trials also produced a more unexpected result: high uptake of crushed bee pupal material. Dyed mite counts were highest for crushed pupae rather than any fungal isolate, while uptake from crushed larvae was markedly lower. This is the opposite of what might be predicted if the bee cuticle, which becomes thicker and potentially more resistant to penetration through development, presents a barrier to feeding. This pattern does correspond though with prior observations of *P. furnarius* behaviour within brood cells, where mites spent most of their time on brood cell walls during the egg and larval stages, but shifted to standing on the bee itself during the pupal stage (Pantoja et al. 2026).

However, feeding behaviour under our trial conditions cannot confirm direct feeding on bee tissue in natural host nest environments. Brood components could not be sterilised before use in feeding trials, leaving open the possibility that dyed mites in these treatments were consuming fungi associated with the pupal material rather than pupal tissue itself. The pupal material was also crushed before presentation, so the experiment did not test whether *P. furnarius* can pierce the exoskeleton of intact bee pupae, as would be required for actual parasitism. Whether *P. furnarius* can breach intact bee cuticle remains unknown, since our piercing assays were unsuccessful. If the mite is unable to penetrate intact bee cuticle, its apparent pupal-stage preference may instead reflect changes in the chemical cues associated with the developing bee, as volatile profiles can change substantially across metamorphosis (Noël et al., 2023). Another possibility is that pupation alters the abundance, composition, or accessibility of fungi associated with the brood, potentially making fungal resources associated with pupae more attractive or accessible to the mites.

*P. furnarius* mouthpart morphology additionally supports a fungivorous or phytophagous feeding mode. Across mites, dietary mode is often reflected in characteristic modifications of the chelicerae and pedipalps (Shatrov, 2012; Di Palma et al., 2009; Nuzzaci & Di Palma, 2002; Di Lillo et al., 2001; Hallas, 1982; Blake et al., 1978; Hislop & Jeppson, 1976). Predatory and parasitic mites typically possess hooked, pincer like, or toothed stylets suited to piercing and gripping prey, alongside clawed or heavily setose pedipalps (Shatrov, 2012; Di Palma et al., 2009; De Lillo et al., 2001; Hallas, 1982; Blake et al., 1978). In contrast, *P. furnarius* had straight, needle-like stylets with only slight distal curving, while its pedipalps lacked claws or coarse setae, more closely matching the morphology of known fungivorous mites (Nuzzaci & Di Palma, 2002; De Lillo et al., 2001; Hislop & Jeppson, 1976). Although this morphology does not exclude feeding on immobile developing bees, which presumably does not require the adaptations used by active predators or ectoparasites, the combination of stylet and palpal features is more consistent with fungivory than predation.

The stylets may also function in probing potential food sources. Chemoreceptors at their tips may provide a mechanism for detecting fungal volatile organic compounds (VOCs) (Günther et al., 2019; Kandasamy et al., 2019; Li et al., 2018; Paludo et al., 2018), as described in other mites (Shatrov, 2012; De Lillo et al., 2001). Fungal VOC profiles vary among species, but many individual compounds are produced across multiple fungal groups and arthropods can respond to structurally related compounds rather than distinct individual compounds (Günther et al., 2019; Kandasamy et al., 2019). This provides a plausible mechanism by which *P. furnarius* could locate multiple fungal food sources sharing common chemical cues, consistent with the range of fungal isolates consumed in the feeding trials and supporting a generalist fungal feeding strategy.

Molecular analysis of the fungal community associated with *P. furnarius* adds a further dimension to this picture. An unexpected observation was a single ASV that dominated the ITS amplicon data across all samples and conditions, including under starvation. Its persistence suggests that this sequence represents a close association with the mite rather than recently ingested food, since a consumed fungus would be expected to fluctuate with availability. Sequence analysis could not confidently assign the ASV to any known fungal taxon: query coverage against its closest BLAST hits was consistently low, top matches spanned taxonomically disparate fungal lineages, and conserved sequence was restricted to the flanking regions while the central region remained divergent from available references.

This pattern is consistent with a taxonomically unresolved fungal associate, but an alternative explanation is that the dominant sequence could derive from host rDNA. Fungal ITS primers can cross-amplify non-fungal sequences, including plant rDNA (Martin & Rygiewicz, 2005), although this has not, to our knowledge, been documented in arthropods. Distinguishing these possibilities is not achievable with amplicon data alone and will require genomic sequencing approaches. If this sequence does prove to represent a fungal associate, its abundance is notable given it has not been detected in ongoing, independent surveys of *T. carbonaria*-associated fungi (unpubl. data). Other mite lineages, including some bee-associated groups, are known to carry and cultivate specific fungi within specialised structures such as sporothecae (Khaustov et al., 2024; Ebermann & Hall, 2004; Moser, 1985), raising the possibility that *P. furnarius* may have a similar adaptation. Whether such a structure is present in *P. furnarius*, however, remains to be determined through detailed morphological examination.

Beyond this putative fungal associate, the remaining fungal taxa identified were each detected at low abundance and only in a single hive. This inconsistent pattern makes it difficult to draw firm conclusions about their ecological significance to *P. furnarius*. *Aspergillus* and *Penicillium* are common saprophytes of pollen and stored food in stingless bee nests (Rutkowski et al., 2023; De Paula et al., 2021) and could reflect transient or opportunistic feeding, or sporadic environmental exposure. *Malasezzia,* however, is a common human- and animal-associated skin fungus (Ianiri et al., 2022) and its presence more likely reflects incidental contamination than a genuine mite association.

## Conclusion

Taken together, these findings suggest that *P. furnarius* is best understood as a facultative fungivore with a flexible association with its bee host. Rather than representing a strictly parasitic or mutualistic interaction, its ecological role may depend on the resources available within the brood cell and the wider colony environment. This flexibility may allow the mite to exploit changing microbial resources within the nest, while also creating the potential for interactions with developing bee brood. Understanding when and why these different feeding strategies occur will be important for determining the ecological significance of *P. furnarius* and its contribution to the broader microbial ecology of stingless bee colonies. More broadly, this study highlights the complexity of mite-bee associations and the value of considering these relationships beyond simple classifications of mutualism or parasitism.

## Statements & Declarations

### Data Availability

Raw amplicon sequences obtained during this study are publicly available in the NCBI Sequence Read Archive under Bioproject ID PRJNA1543362. All other data that support the findings of this study are available in the supporting material of this article.

## Acknowledgements

We thank Julianne Lim for her technical assistance and Owen Seeman and Matt Shaw for their assistance interpreting mite morphology. We acknowledge the technical and scientific assistance of Sydney Microscopy & Microanalysis, the University of Sydney node of Microscopy Australia.

## Funding

This work was supported by Australian Research Council Discovery Early Career Award to KEF (DE250100611).

## Competing Interests

The authors have no relevant financial or non-financial interests to disclose.

## Author Contributions

CP, RG, and KEF conceived and designed the experiments. CP, AFP, RS, and RG collected the samples. CP generated the data with contributions from JL. CP collated and analysed the data and wrote the manuscript with contributions from RG and KEF. All authors read and approved the final manuscript.

**Supplementary Table S1.** Identified isolates cultured from Tetragonula carbonaria brood cells, including those used in feeding trials, identified via Sanger sequencing and NCBI BLASTn.

**Supplementary Table S2.** Selected top BLAST hits for the dominant unclassified fungal ASV (ASV1), following primer trimming.

**Supplementary Table S3.** Summary of different colony morphological types grown on agar plates from brood cells.1

## Notes

### Competing Interest Statement

The authors have declared no competing interest.

